# *Z*-score to evaluate the absolute quality of macromolecular models based on SAXS data

**DOI:** 10.1101/349704

**Authors:** Dmytro Guzenko, Sergei V. Strelkov

**Affiliations:** Department of Pharmaceutical and Pharmacological Sciences,KU Leuven, Leuven 3000, Belgium.

## Abstract

Small-angle X-ray scattering (SAXS) is a highly popular technique to assess the native three-dimensional structure of biological macromolecules in solution. Here we introduce a statistical criterion, Z-score, as a novel quality measure of SAXS-based structural models which positively correlates with data quality. We propose that, besides a goodness-of-fit (GOF) measure such as reduced χ^2^, the Z-score reflecting the ability of a given SAXS curve to differentiate between possible models should always be reported.

---

The main benefit of SAXS is its ability to rapidly characterize the structure of biomacromolecules and complexes thereof in a native solution environment [1], [2], [3]. Recent progress in hardware as well as in data processing algorithms have resulted in an explosive growth of the technique’s popularity [4]. Since the solute particles in a typical biological SAXS experiment are randomly oriented, the resulting SAXS signal is a one-dimensional profile *I*(*q*) which corresponds to a spherical average of the particle’s Fourier transform. This averaging leads to a major information loss. As a result, for typical biological molecules, determination of the 3D structure from SAXS data represents an ill-posed problem [1]. Correspondingly, reliable *ab initio* shape reconstruction even at a modest resolution necessarily requires additional constraints [5].

At the same time, if one or more ‘test’ 3D models are available then the expected SAXS signal can be calculated from theory and subsequently scaled to the experimental data [6]. The quality of the resulting fit is usually assessed by the reduced χ^2^ [7], calculated over ~ 10^3^ data points of the *I*(*q*) profile. A χ^2^ value close to 1 is taken to signify a successful modelling. This practice, however, stumbles upon an intrinsic limitation, namely that the χ^2^ only reflects the quality of the model relative to the *given* dataset [8]. In particular, noisy experimental SAXS curves naturally produce lower χ^2^ values, which may even encourage non-expert users to rely on inferior data. A recently proposed GOF test, CorMap, while not requiring explicit experimental error estimates, does not resolve this limitation [9]. Moreover, the reliability of several structural models of the same biomolecule based on different SAXS datasets cannot be directly compared. This situation occurs, for instance, when conflicting SAXS-based models for the same system are proposed by different research groups.

We reasoned that the significance of a structural model can be assessed upon analysing the discriminative power of the underlying SAXS curve with respect to a continuum of all models possible. To the latter end, the Protein Data Bank (PDB) can provide a good reference, as it covers practically all plausible sizes and shapes of biomacromolecules and their complexes [10], and can therefore represent the variability of the expected SAXS signal. Consequently, we have developed a specific statistical measure, *Z*-score, which indicates how the fit between the theoretical scattering from the current model and the experimental dataset stands out from a sample of equivalent fits calculated for a representative sample of PDB structures. The use of *Z*-scores has proven instrumental in various fields, including macromolecular crystallography [11], [12].

Towards a procedure to evaluate the SAXS-specific *Z*-score, we have first prepared a reference collection of scattering curves from known macromolecular structures. Briefly, scattering curves were calculated for a non-redundant collection of ~23000 PDB entries, and thereafter ~4000 curves were selected to uniformly sample the space of all possible SAXS curves (See Methods and Fig. S1). The *Z*-score for a structural model against a given experimental dataset can then be calculated as follows. Initially all curves from the reference set are fitted to the experimental curve (Fig. 1a). The distribution of the resulting χ^2^ values, normalized to the maximal value 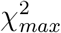, is produced and approximated by a beta distribution (See Methods and Fig. S2). Subsequently, a fraction of the reference models that have χ^2^ values not exceeding 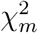 of the given structural model is estimated (Fig. 1b). This fraction effectively corresponds to the probability of the current model yielding the 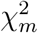 value by a mere chance. An established way of expressing this probability is through the so-called *Z*-score of a standard normal distribution [13]. Such score conveniently describes the ‘uniqueness’ of theoretical scattering from the model being tested with respect to the given experimental curve. Here, obtaining a high *Z*-score implies that a randomly picked reference curve is very unlikely to produce a fit with 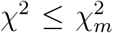 whereas a *Z*-score of 0 corresponds to a 50:50 chance.

**Figure 1.**
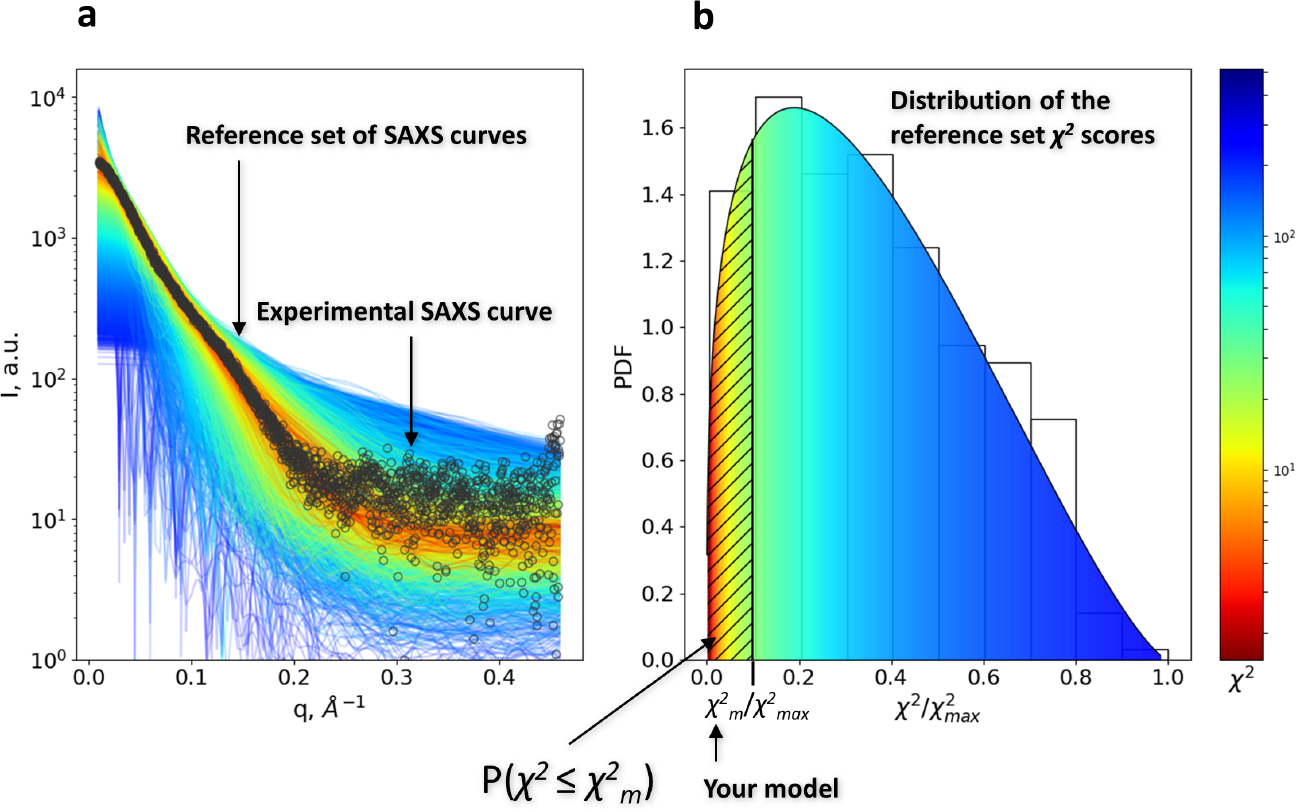
The principle of Z-score calculation. (a) The reference set of 4000 SAXS curves fitted to an experimental SAXS curve (SASBDB entry SASDAM5, black circles). The reference curves are coloured according to their χ^2^ value with respect to the experimental curve. (b) Probability distribution for the 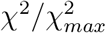 ratio. The proportion of models from the reference set with χ^2^ not exceeding that of a given model is represented by a hashed area.

The same approach immediately allows us to introduce an absolute quality measure of an experimental dataset, *Z*^0^, independent of a particular model. We define it as the *Z*-score of a statistically sound ‘perfect’ model, which can be readily obtained *via* the procedure illustrated in Fig. 1b for 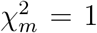. The *Z*^0^ value describes a dataset quality in terms of its discriminatory power with regard to any theoretically possible profile. It also corresponds to the upper limit of the *Z*-scores that can be obtained upon refining a structural model. Correspondingly, the ratio *Z/Z*^0^ can serve as an efficient criterion of the convergence between the model and the data.

Importantly, in contrast to the *χ*^2^ value, the *Z*-score positively correlates with the quality of underlying SAXS data. This can be illustrated by a solution SAXS analysis of the α-crystallin domain dimer of human small heat shock protein HSPB1. While several distinct dimeric arrangements can be considered (Fig. 2), only one of them (denoted as AP_II_) was confirmed though chemical crosslinking and other methods [14]. As can be seen, a relatively noisy experimental curve with *Z*^0^ = 1.6 yields a reasonable χ^2^ value of 1.4 with respect to the non-native AP_III_ dimer. However, the χ^2^ value of 2.9 obtained upon fitting the correct dimer AP_II_ to the higher-quality experimental curve with *Z*^0^ = 2.43 would be considered unsatisfactory. In contrast, the *Z*-scores give a clear preference to the correct model. Notably, CORMAP [9] rejects both data-model fits shown on Fig. 2 with equal P-values of 0. At the same time, elastic deformation of the model using the popular SREFLEX [15] program yields a χ^2^ ≃ 1 for either data-model pair. Thus both the correct AP_II_ and the wrong AP_III_ dimer are judged as equally (in)significant by the current tools.

**Figure 2:**
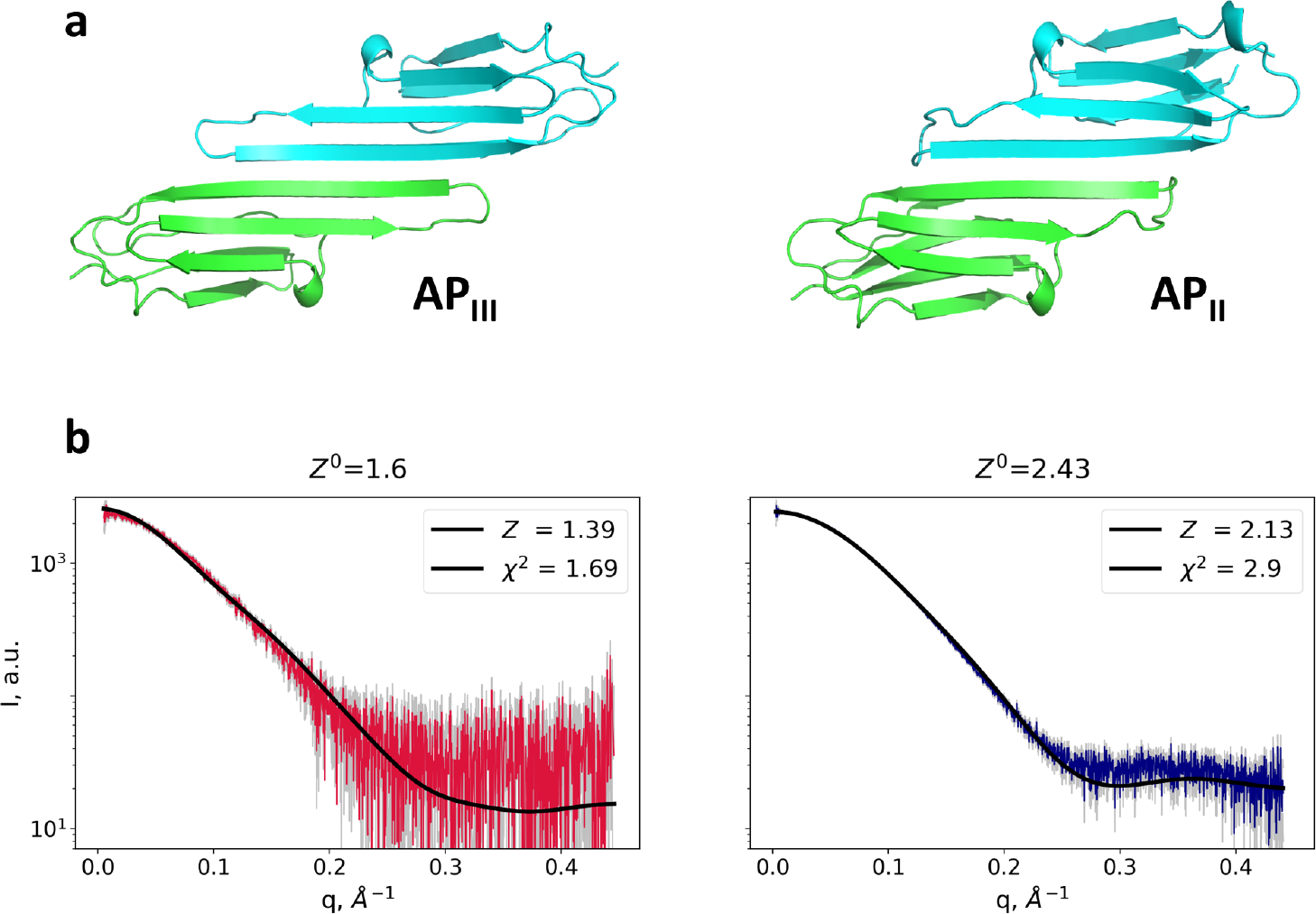
Evaluation of the model-data fits by χ^2^ and *Z*-score. (a) Two possible atomic models of the α-crystallin domain dimers of human small heat-shock protein HSPB1 shown as ribbon diagrams [14]. (b) Theoretical scattering (black curves) of the AP_III_ and AP_II_ models were fitted to the experimental datasets collected with a diluted (2 mg/ml, red curve) and concentrated (11 mg/ml, blue curve) protein samples respectively. The resulting *Z*^0^, χ^2^ and *Z*-scores are indicated on the plots.

To explore the relationship between the *Z*-score and the root mean square deviation (RMSD) of a test model with respect to the true structure we have used a simulated example. Atomic structure of transportin-SR2 extracted from the PDB entry 4OL0 was taken to represent the true solution conformation. Four simulated SAXS curves were produced by calculating the scattering from this structure and supplying experimental noise at four different levels, and a family of ~100 test models was created upon an elastic deformation of the crystal structure [16] (see Methods). The result of fitting the theoretical scattering from every model to each of the four simulated SAXS curves is presented in Fig. 3. Notably, the noisier curve (*Z*^0^ = 2.89) can be fitted using numerous models with a χ^2^ close to 1, many of which greatly deviate from the crystal structure (RMSD > 10Å). In this case, low χ^2^ values alone do not warrant a structural agreement between the test model and the true one. In contrast, only a few test structures fit the most accurate curve (*Z*^0^ = 4.15) with χ^2^ below 1.5, and all of them are reasonably close to the true structure (RMSD < 5Å). Hence, only the experimental data with the highest *Z*^0^ value provide for a convergence of modelling to the true structure.

**Figure 3:**
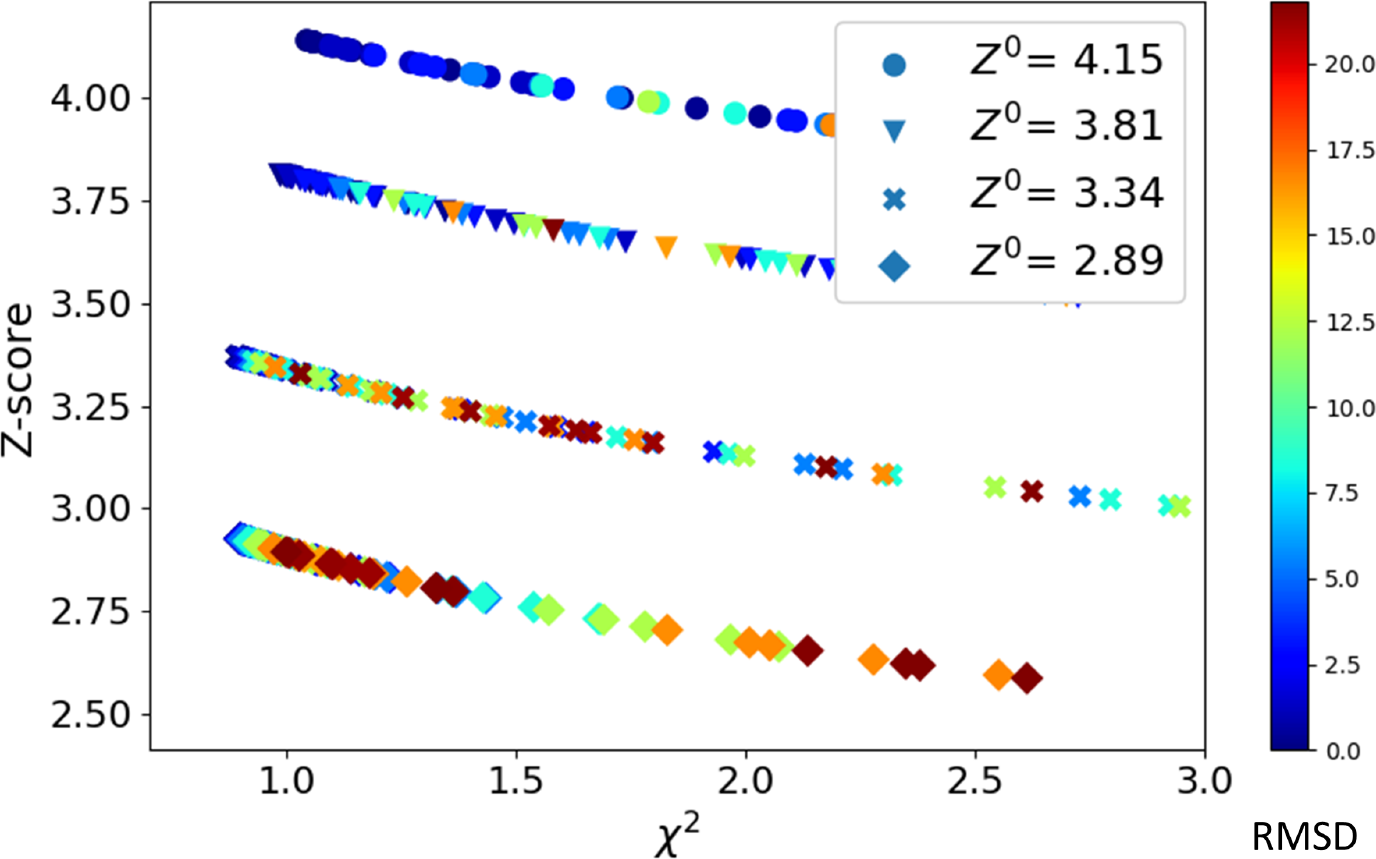
Relationship between *Z*-score, χ^2^ and the structural similarity between test models and the true structure, illustrated by a simulated example. For four simulated SAXS profiles with *Z*^0^ as indicated and a series of test models, *Z*-scores (vertical axis), the corresponding χ^2^ (horizontal axis) and the RMSD relative to the true structure (rainbow colouring) are plotted.

We have also surveyed the biological SAXS profiles from the SASBDB database [17] (~400 datasets at the time of writing). As can be seen in Fig. S3, their *Z*^0^ values range from 0.99 to 7.18 (median 3.17). Interestingly, while the *Z*^0^ values tend to correlate with the size of the particles as reflected by their maximal dimension *D*_*max*_, there is a wide spread of *Z*^0^ even for particles of comparable size. This observation suggests that the data quality (which depends of the quality of experimental setup, protein concentration, radiation exposure, *etc.*) makes a major contribution to the *Z*^0^ value.

In summary, the *Z*-score introduced here appears an extremely useful addition to the current practice of SAXS-based structural modelling. We suggest that for each model obtained both the χ^2^, as the most commonly used GOF measure, as well as the *Z*-score should be reported. χ^2^ ≫ 1 unambiguously indicates a failed modelling, which can be due to major flaws of the experiment, wrong assumption on the sample composition, or an excessively restrained 3D shape. If, however, a χ^2^ ≈ 1 is achieved, the corresponding *Z*-score can indicate the absolute significance of the obtained model.

The *Z*^0^ score provides a ‘probabilistic resolution’ of an experimental SAXS curve, reflecting the amount of structural information contained in the curve given its noise level. Of note, a recently proposed algorithm Ambimeter [18] represents inherent ambiguity of a SAXS profile through a number of distinct 3D shapes that it can account for. This is expressed by a so-called a-score which only depends on the shape of the solute particle (which can not be changed). Importantly, our *Z*^0^ score, which critically depends on the quality of the experimental data (which can be improved), and the a-score of Ambimeter essentially complement each other, jointly characterizing a potency of a given SAXS curve towards producing a reliable 3D model. Indeed, it should be expected that cases corresponding to higher ambiguity scores should require data with higher *Z*^0^ values to enable the convergence of structural modelling to a unique solution (such as in example in Fig. 3). Further research along these lines should help approaching the central challenge of SAXS-based modelling of biomacromolecules, namely a reliable quantification of the level of structural detail that can be achieved through a given SAXS experiment.

## Methods

Methods and associated figures and references are available in the online supplement.

## Acknowledgements

The authors thank all colleagues in their lab and beyond who contributed to the discussions on SAXS data evaluation and the use of the *Z*-score metric. This research originated from a number of biological SAXS-based projects (to S.V.S.) funded by the FWO (Research Foundation - Flanders) and KU Leuven. SAXS data collection was made possible through the SWING beamline at Synchrotron Soleil and the P13 beamline of DESY.

